# Adaptation and trade-offs with insecticide resistance in overwintering Drosophila

**DOI:** 10.1101/2025.05.27.655643

**Authors:** Eric G. Prileson, Bianca Campagnari, Catherine I. Clare, Amir R. Gabidulin, René S. Shahmohamadloo, Seth M. Rudman

**Affiliations:** Washington State University Vancouver, School of Biological Sciences,14204 NE Salmon Creek Ave. Vancouver, WA, USA 98686

## Abstract

Winter is a formidable challenge for ectotherms that inhabit temperate climates. Prior work has demonstrated that multivoltine organisms can evolve rapidly in response to temporal changes within a growing season, a process termed adaptive tracking. However, the mechanisms by which winter conditions drive rapid adaptation, particularly when combined with strong anthropogenic stressors, remain poorly understood. Here we use replicate populations of *Drosophila melanogaster* in a field experiment to test i) whether winter conditions drive rapid adaptation and ii) for trade-offs between insecticide resistance and overwintering survival. Following a longitudinal field experiment spanning summer and fall investigating the evolution of insecticide resistance, we tracked subsequent evolution during an overwintering period. We detected repeated evolutionary shifts indicative of adaptation to winter conditions in multiple traits, including body size and fecundity. Additionally, populations that had evolved insecticide resistance during the growing season had reduced survival and showed patterns of lower resistance following the winter period, suggestive of a trade-off between overwintering success and insecticide resistance. These rapid evolutionary responses and potential trade-offs provide important context for understanding overwintering performance in temperate insects, with implications for pest management and ecosystem services.

## Introduction

Rapid adaptation is critical to organismal responses to environmental change and the maintenance of biodiversity (Exposito-Alonso et al. 2022; Urban et al. 2016). There has been considerable work to understand the ecological drivers of rapid adaptation, including biotic drivers such as predation, competition, and mutualism (Li et al. 2021; Reznick and Endler, 1981; Schluter, 2000). Similarly, abiotic factors, including variation in temperature, have been studied as drivers of rapid adaptation with direct implications for responses to climate change (Bradshaw and Holzhapfl, 2010; Huey et al. 2021; Radchuk et al. 2019). In ectotherms, there is extensive study on the ecology and physiological challenges associated with winter (Denlinger and Lee, 2010; Marchand, 1987; Sinclair et al. 2003), but compared to summer, the role of winter conditions in driving rapid adaptation has received comparatively little attention (but see Campbell-Staton et al. 2017, Marshall et al. 2020, and Williams et al. 2014). Further, there are even fewer studies having sufficient population-level replication required to detect parallel changes indicative of adaptation to winter conditions. Although there is clear evidence of ecological and demographic effects of winter conditions, including population declines and life-history trade-offs between winter stress tolerance and summer reproduction (Boulétreau-merle and Fouillet, 2002; Marshall et al. 2009), there is little known about how overwintering selection shapes evolutionary trajectories.

The fate of overwintering ectotherm populations is largely dependent on both physiology and demography. The physiological mechanisms enabling ectotherms to survive challenging winter conditions are well-documented, including the production of cryoprotectants, expression of antifreeze proteins, and maintenance of cell membrane fluidity and ion balance (Sinclair et al. 2003; Teets et al. 2023; Toxopeus and Sinclair, 2018). Further, both thermoregulatory behavior (e.g., seeking thermal refugia; Denlinger and Lee, 2010; Hoffman et al. 2003; Sinclair et al. 2003), and life history strategies including facultative and reproductive diapause (Schmidt and Conde, 2006) can increase overwinter survival but may be associated with reduced fecundity (Meuti et al. 2024; Erickson et al. 2020; Schmidt and Conde, 2006). In addition to putatively adaptive physiological responses, for multivoltine species, population size and number of generations per year are strongly influenced by seasonal fluctuations in temperature (Altermatt, 2010; Roff, 1980; Tauber and Tauber, 1981; Shpak et al. 2010). These demographic shifts are important for ecological interactions (Markow and O’Grady, 2008; Parmesan, 2006; Thackeray et al. 2016) but also for the capacity for rapid adaptation, because they alter genetic diversity of populations (Barrett and Schluter, 2008; Crozier and Dwyer, 2006). Across winter, these populations can be subject to severe bottlenecks (Chen et al. 2006; Kinnison et al. 2007; Lawton et al. 2022) where demographic effects of genetic drift and frequency of adaptive physiological traits likely play a role in overwintering persistence. The evolutionary implications of winter population collapse and expansion the following spring, however, remain poorly understood.

There is evidence of physiological responses to co-occurrence of low temperature and natural stressors such as desiccation or pathogen exposure (reviewed in Sinclair et al., 2013; Zhang et al., 2011; Le Bourg et al., 2009). However, how the emergence of novel, strong, directional selective agents – such as those imposed anthropogenically – interact with natural stressors to shape ecological and evolutionary outcomes (e.g., insecticide resistance and low temperature) are less clear (ffrench-Constant and Bass, 2017; Kliot and Ghanim, 2012). For example, there could be positive correlational selection and genetic covariance between insecticide resistance and overwintering survival. Here, selection and heritability for one response is positively associated with the other (Lande and Arnold, 1983; Service and Rose, 1985; Sinervo and Svensson, 2001). Alternatively, there could be antagonism – negative correlational selection and genetic covariance – of insecticide resistance and overwintering. Here, resistant populations are less likely to survive the overwintering period (McKenzie, 1990; McKenzie, 1994), possibly due to pleiotropic effects and/or potential fitness trade-offs (Crow, 1957; Roush and McKenzie, 1987). These outcomes for resistance across seasons are important considerations for agriculture as there is a need to understand the mechanisms that facilitate or constrain insecticide resistance evolution (Sparks et al. 2012; Baker et al. 2007) and allow more efficient use of insecticides as part of broader pest management strategies.

The common fruit fly, *Drosophila melanogaster,* makes an excellent system to study natural selection across winter. In addition to the exceptional molecular and population genetic tools and amenability as a model organism, *D. melanogaster* is among the species most commonly studied in an overwintering context including responses to low temperature (Hoffman, 2003; Overgaard et al. 2014; MacMillan and Sinclair, 2011; Rako and Hoffman, 2006; Schmidt et al. 2005). Despite a tropical ancestral origin, natural populations of *D. melanogaster* are cosmopolitan and can overwinter even at high latitudes. Although it is generally suspected that high-latitude populations take advantage of human dwellings or compost piles as overwintering refugia (Ives, 1970), *D. melanogaster* does have recently derived physiological and life history traits for maintaining homeostasis under freezing conditions. For example, female flies undergo a reproductive diapause, produce cryoprotectants, and can maintain membrane fluidity and ion gradients at low temperatures (Denlinger and Lee, 2010; Hoffman et al. 2003; Sinclair, 2003). Despite these adaptations, mortality in winter can reach 90% or greater, though field survival data is scarce (Ives, 1970; Izquierdo, 1991; Mitrovski and Hoffman, 2001; Nuñez et al. 2024). Evidence from low temperature responses of flies, however, do show patterns of latitudinal variation in putative winter adaptations. At the same time, spatial patterns of genomic variation suggest that high latitude populations exhibit genomic patterns of local adaptation with relatively high effective population sizes to sustain populations overwinter (Denlinger and Lee, 2010; Ives et al. 1970; Collett and Jarman, 2001; Cogni et al. 2015; Machado et al. 2021). Thus, it is uncertain how evolution across winter allows *D. melanogaster* to maintain genetic variation and expand successfully in spring despite population bottlenecks.

Temperate populations of *D. melanogaster* can adapt in response to temporal environmental fluctuations that shift within and among generations – a process termed adaptive tracking (Bergland et al. 2014; Rudman et al. 2022; Behrman et al. 2015). As such, prior evolution to outdoor conditions might shape the evolutionary trajectory of overwintering populations (Grainger et al. 2022) with certain directions expected in response to winter selection. Adaptation could include reduced fecundity due to higher reproductive diapause incidence (Collet and Jarman, 2007; Schmidt et al., 2005), greater stress tolerance (e.g., improved cold and starvation tolerance; Hoffman, 2010; MacMillan and Sinclair, 2011) due to life-history allocation trade-offs (Stearns, 1998), and larger body size from the “temperature-size rule” (Angilletta, 2009; Chown and Nicholson, 2004; Partridge, 1994). Fall fly populations have exhibited phenotypic and genomic signatures associated with higher thermal tolerance but worse cold tolerance (i.e., more ‘summer like’; Bergland et al., 2014; Behrman et al., 2015). Given that insecticide treatments are not commonly applied in winter, insecticide-resistant fall populations that overwinter as adults would face selective pressure from low temperatures while likely incurring costs for carrying resistance (Roush and McKenzie, 1987; McKenzie, 1994). This could mean that resistant alleles decline in frequency across winter, an important factor for pest population control measures.

Here, we use a field experiment to assess both the magnitude of overwintering adaptation and the potential for trade-offs or covariation with adaptation of insecticide resistance. Specifically we ask the following questions: 1) Does an overwintering period drive adaptation as measured by parallel genetic change across populations? 2) Is there a trade-off between selection during the overwintering period and prior selection for insecticide resistance? 3) Does adaptive tracking across a summer and fall growing season confer fitness benefits for overwinter survival? We hypothesized that strong overwinter mortality drives adaptation for greater overwintering performance (Izquierdo, 1991; Mitrovski and Hoffman, 2002). Therefore, we predicted that flies that successfully overwinter would exhibit greater body size, reduced fecundity, and greater starvation tolerance than their fall counterparts. For our second question, we define a trade-off as the observed negative association between phenotypic measures including survival; a broader definition of the term ‘trade-off’ that describes the outcomes from rather than the mechanistic causes of a trade-off (Garland et al. 2022). We expected that overwintering survival would negatively covary with resistance to an insecticide (McKenzie, 1994; Miyo, 2000). We predicted that overwintering selection would reduce insecticide resistance and, in turn, that resistant populations would have lower survival and reduced performance of cold tolerance traits. Lastly, we hypothesized that temporal adaptation to the outdoor environment is adaptive for overwintering and would also produce wider variation in responses than populations that had not experienced seasonal selection.

Assessment of rapid evolutionary adaptation and potential tradeoffs requires replicate populations that undergo selection in parallel. However, such studies are often carried out in highly controlled laboratory conditions (but see Hoffman et al. 2003; Sgrò and Hoffman, 1998). Replicated field experiments allow for environmental realism but retain the replication needed to connect patterns to processes. This is particularly important in the context of winter where both average and extreme low temperatures of winter cause physiological thresholds to be crossed repeatedly, leading to fitness impacts (Overgaard and MacMillan, 2017; Williams et al. 2015; Marshall et al. 2020). In this vein, we established *D. melanogaster* populations into 40 replicated outdoor mesocosms in the summer and tested for parallel phenotypic evolution across independent populations. With a subset of these original populations, we then used repeated common garden rearing following an overwintering period to determine whether overwintering drove adaptation and if there was negative covariance of resistance and overwintering performance. By testing for parallel changes across independent overwintering populations, we examine evidence for adaptation across winter, interactions with evolved insecticide resistance, and provide important context to the well-established physiological mechanisms of overwintering.

## Materials and Methods

### Multigenerational experiment and fly populations

We tested fly populations as part of a multigenerational selection experiment following previously described protocols outlined in Rudman et al. (2022). *D. melanogaster* populations were founded from 100 DGRP laboratory fly lines (Drosophila Genetic Reference Panel; Mackay et al. 2012) recombined in the lab for nine generations prior to outdoor rearing. Populations were reared for approximately nine generations in outdoor mesocosms located 45.729 N, –122.633 W (from here on “orchard”) from July 17th until October 30th, 2023. Throughout the experiment, all populations were fed a modified Bloomington recipe media.

Fly populations included those fed control media (from here on “control” populations) and populations fed exclusively media treated with a 0.0375 ug L^-1^ concentration of the organic insecticide spinosad [Entrust; Dow AgroSciences, Indianapolis, IN, USA]. This concentration corresponds with the lethal dose at which 50% of flies saw mortality from in-house trials of egg-to-adult survivorship. Spinosad is a widely used insecticide that targets the Dα6 subunit of nicotinic acetylcholine neuronal membrane receptors (nAChR), leading to neuronal overexcitation and ultimately mortality in larvae (Martelli et al. 2022; Salgado, 1998; Perry et al. 2007). Of the original insecticide-exposed populations, half of the cages went extinct during the growing season with the remainder persisting – possibly via evolutionary rescue through the evolution of spinosad resistance – from standing genetic variation (Perry et al. 2007; Perry et al. 2021). From here on we refer to the insecticide-exposed populations that evolved resistance during the growing season as “resistant” populations. A third subset of outbred founder populations (from here on “founder” populations) were maintained in the laboratory on control media at 25 °C with a 12:12 LD cycle at moderate densities and were not exposed to outdoor conditions across the growing season (Figure 1).

**Figure 1.**
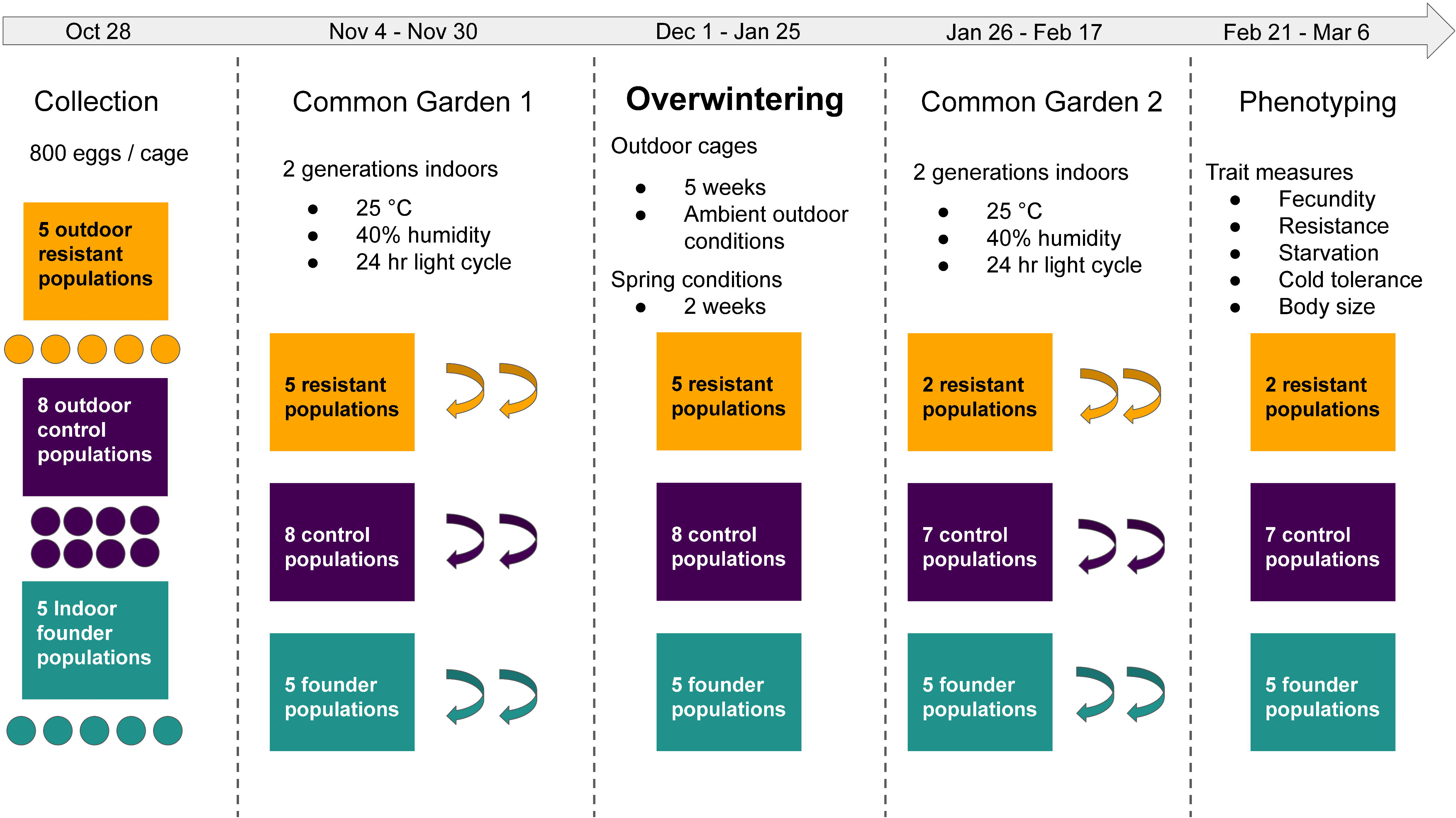
Schematic of the overwintering experimental design. Colors indicate population types of flies (see main text for details), shapes indicate independent replicate populations, and colored arrows indicate one generation of laboratory common garden rearing. Dates are for the winter of 2023 and 2024.

### Collection and initial common garden rearing

At the end of the growing season (October 28, 2023; Figure 1), we collected approximately 800 eggs per cage in density controlled 200 mL plastic bottles from eight outdoor control, five outdoor resistant, and five indoor founder populations. We then reared flies indoors under common garden conditions at 25 °C under 12:12 L:D lighting and 40% humidity for two generations to control for density effects among outdoor populations. Upon eclosion of the second generation reared under common conditions, we collected approximately 2000 adult flies per population for the overwintering experiment.

### Experimental treatments and overwintering trial

The orchard experiences a relatively mild temperate winter typical of the western Cascades region of the Pacific Northwest, but it still presents challenging conditions for *D. melanogaster* including occasional sub-freezing air temperatures (Figure S1, S2). To provide a favorable environment, we used 5L clear plastic containers with a small, screened opening to allow aeration and added layers of hay pellets, ash wood chips, and cotton for insulation. We acknowledge that these do not provide endogenous heat sources and are likely more challenging than conditions where flies are hypothesized to overwinter (e.g., agricultural compost piles; Ives, 1970). We then transferred 2000 flies from each independent population and each treatment type (control, resistant, founder) into an overwintering container along with two 50 mL control (i.e., non-spinosad treated) food media bottles as a food resource. We placed the overwintering flies first into an outdoor shed for 24 hours to acclimate to low (but less extreme) temperatures prior to transfer to a large 2 m^3^ outdoor rearing cage.

After five weeks in the outdoor overwintering trial, we transferred the flies to a greenhouse to simulate a spring-like phenology cue in both temperature and photoperiod. The greenhouse was held with temperatures at 15 °C with a 16:8 light dark cycle and we added a fresh food bottle with yeast paste. After one week in the greenhouse, we then transferred flies to an indoor temperature controlled incubator (Percival DR-41 NL) where we increased the temperature cycle daily to simulate spring phenology cues and encourage reproduction (Table S1).

During the overwintering period and spring acclimation process, four populations (three resistant and one control) saw complete mortality with zero reproduction and so we removed these populations from the remainder of the experiment. This left a final sample size of two resistant, seven control, and five founder populations and we counted the total number of surviving adults from each population. We then collected eggs from the overwintering flies in four media bottles per cage at a density of 200 eggs / bottle and moved these media bottles to a common garden rearing room kept at 25 °C, 12:12 LD and 40% humidity. We expanded post-overwintering flies for two generations and three to five days after eclosion, we began our phenotyping trials (Table S2).

### Phenotypic assays

To assess temporal evolutionary responses during the growing season, eggs were collected from each replicate mesocosm at regular intervals (August, September and October) and reared in common garden conditions as described above. The following phenotypes were assayed per replicate mesocosm: 1) Insecticide resistance measured as survivorship to adulthood: the proportion of eggs (30 eggs per vial) that survived to adulthood in 3 replicate vials; 2) Fecundity: the total eggs laid by five females over three days, measured in each of 3 replicate bottles; 3) Starvation tolerance: the time to starvation for three replicate vials containing 10 males each on agar-only media; 4) Adult body size, measured as the average dry mass of three pools of five females, dried at 55°C for 24 h. We repeated these phenotypic assays for the overwintering flies but also included a fifth measure: chill coma recovery time (CCRT), a static measure of cold tolerance widely used in assessing low temperature responses in *D. melanogaster* (Gibert and Huey, 2001). We used a chill coma temperature of 0 °C for two hours following Macdonald (2004) and Rako and Hoffman (2006) with 15 female flies in three replicates per cage and scored recovery as the time when flies were able to right themselves.

### Statistical analyses

To test adaptation across winter, we compared phenotypic measures from the populations before fall collection (from here on ‘fall’) and following the overwintering period. We modeled each phenotypic measure as the response variable, time point as our fixed effect predictor variable, and considered cage (i.e., individual population) as a random effect. To assess a potential trade-off in resistance, we tested for differences in survival and evolutionary divergence in traits across time points between outdoor-reared resistant and control populations. Here, each phenotypic measure was the response variable, time point and population type were treated as fixed effects along with their interaction, and cage was treated as a random effect. Since CCRT was only measured in overwintering populations, we tested cold tolerance between overwintered control and resistant populations in a model with population type as the predictor variable and cage as a random effect. Lastly, we tested for evolved differences between outdoor reared control and indoor reared founder populations to test for an effect of seasonal evolution on overwintering success. For this third question, phenotypic measures were response variables, population type was our fixed effect predictor, and cage was a random effect.

To model each phenotypic measure for each question, we used generalized linear mixed-effects models constructed with the *glmmTMB* function from the *glmmTMB* package (Brooks et al., 2017) and applied appropriate link functions to match error distributions (see supplemental information for details on each model). Model diagnostics were conducted using the *simulate Residuals* and *test Dispersion* functions from the *DHARMa* package (Hartig and Lohse, 2022). Some of our models were unable to meet assumptions of homoscedasticity, likely due to the reduced sample size of overwintered resistant populations. Efforts to correct for this did not improve model fits and so we opted to retain original model designs but deemphasized the focus on significance relative to effect size (see supplemental information). We determined significance of the predictors using log-likelihood ratio tests in the *Anova* function with the type = “III” argument, tested equality of variance using the *leveneTest* function from the *car* package (Fox and Weisberg, 2022) and calculated effect sizes as Hedge’s G using the *hedges_g* function from the *effectsize* package (Ben-Shachar et al. 2020). All analyses were conducted in R version 4.3.3 (R Core Team, 2024) with results reported to three significant digits.

## Results

### Field survival of overwintering flies

In our field test of overwintering for the three population types of *D. melanogaster* (control, resistant, and founder), all populations saw severe mortality of >98% from a starting population of 2000 adult flies (Figure 2) with 1.67% of control, 0.84% of founder, and 0.62% of resistant flies surviving. These differences in survival among population types were broadly explained by the complete mortality of three of the five outdoor resistant populations while only one of the eight outdoor control and none of the five indoor founder populations had complete mortality. The model of average overwintering survival among populations showed a non-significant trend of population type on survival (Χ^2^ = 5.35, P = 0.0689).

**Figure 2.**
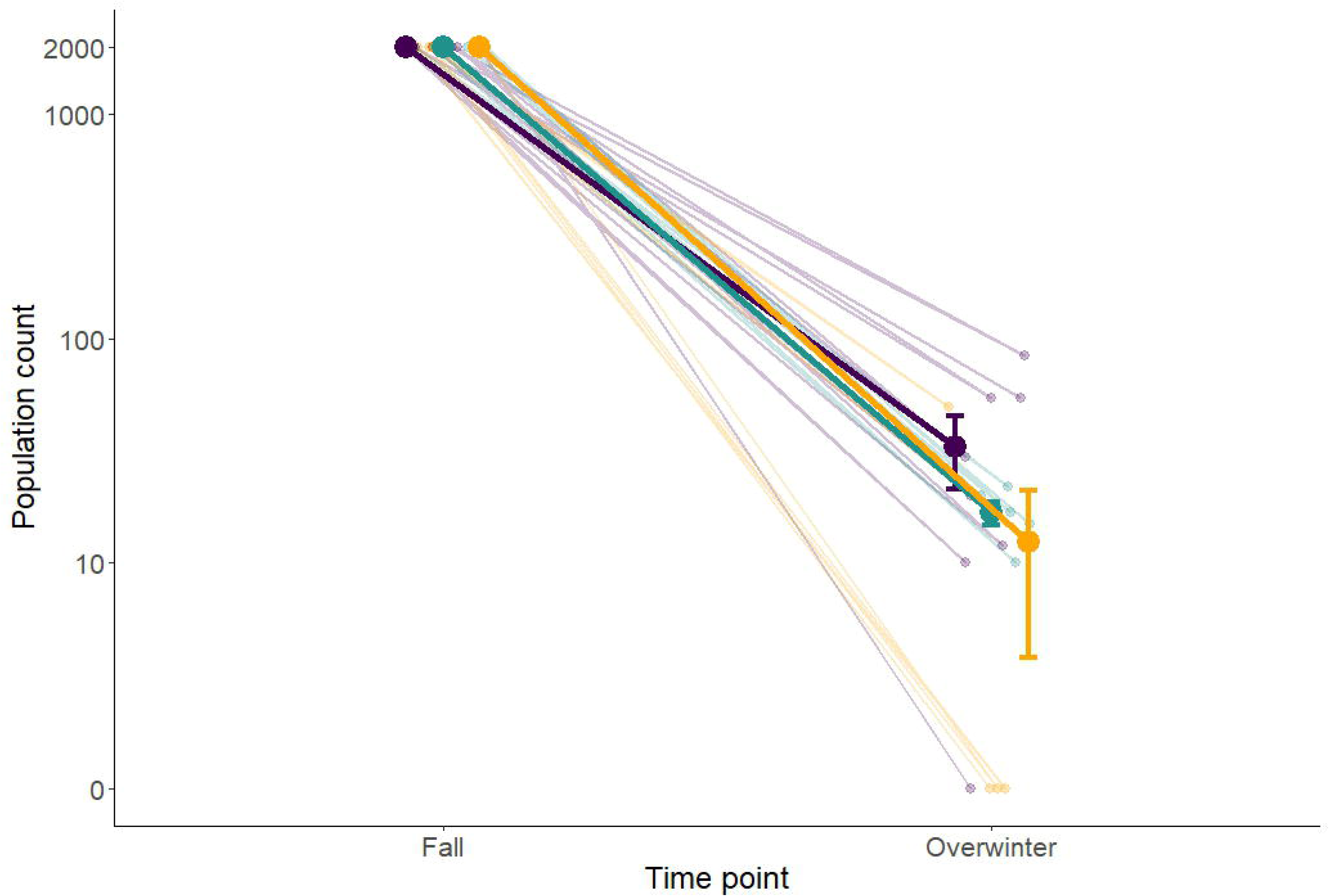
Population counts of flies before and after the overwintering period. The population count on the y axis is on a ‘pseudo’ natural log scale that can accommodate the zero values that indicate population mortality. Large points show the mean number of surviving flies per population type ± 1 SE with colors corresponding to each population type (purple = control cages, green = founder cages, orange = resistant cages; note: SE bars for the founder cages are within the size of the point). Smaller points jittered behind the means show the average number of flies per replicate cage while lines connecting each point show the decline in population size across all replicates.

### Adaptation across winter

We found evidence for parallel temporal evolution in most phenotypes. Compared to fall populations, overwintering populations exhibited smaller dry mass (Figure 3A; estimate = – 0.398, SE = ± 0.0606; Χ^2^ = 43.2, P < 0.0001), greater fecundity (Figure 3B; estimate = 0.515, SE = ± 0.0543; Χ^2^ = 136.7, P < 0.0001), and decreased spinosad resistance measured as percent survival (Figure 3D; estimate = –24.9, SE = ± 2.08; Χ^2^ = 174.7, P < 0.0001). There was no difference in starvation tolerance across the overwintering period (Figure 3C; estimate < 0.0001; SE = ± 1.91; Χ^2^ = 0.00, P = 1.000).

**Figure 3.**
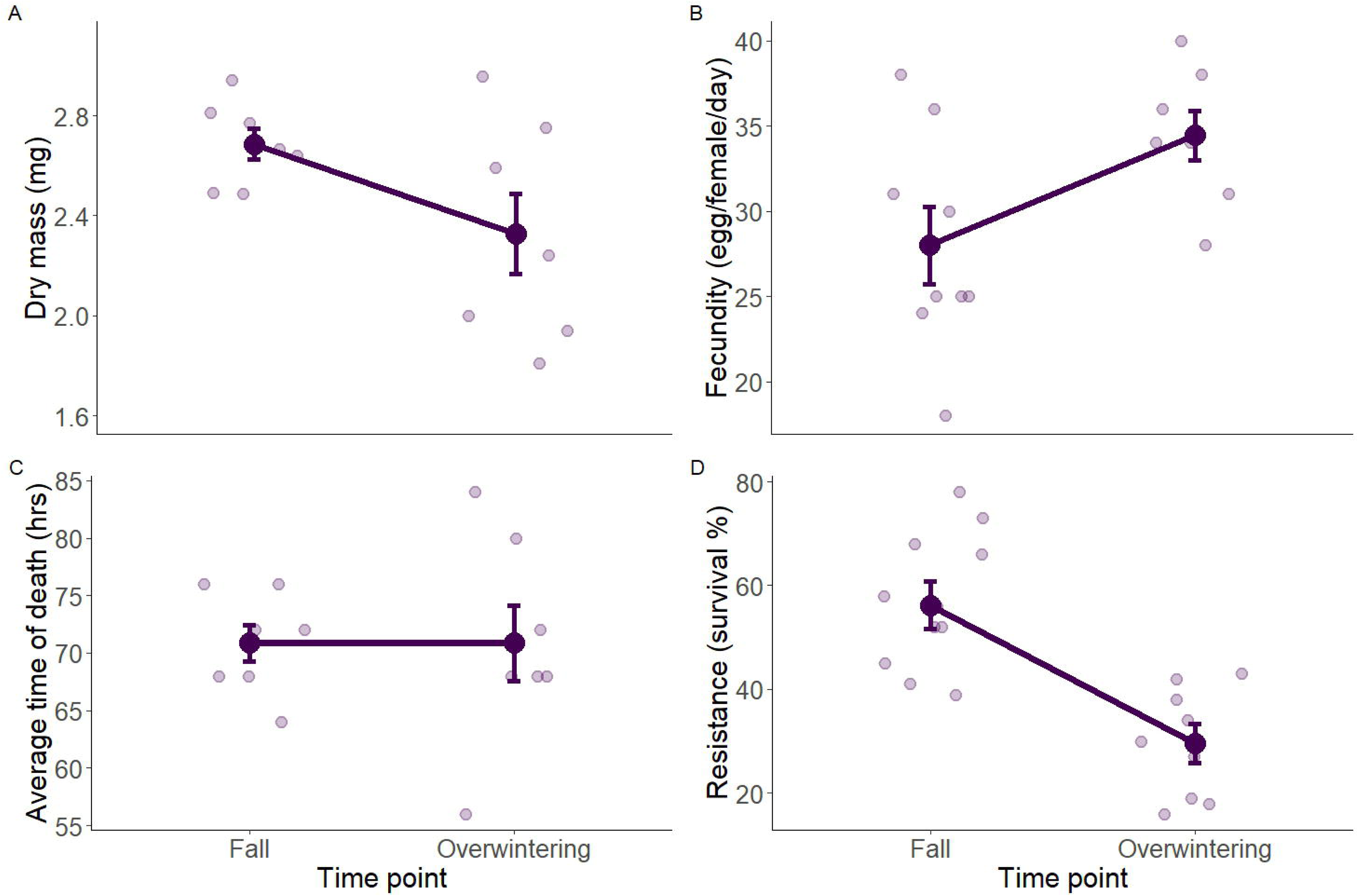
Phenotypic measures of outdoor control populations (control) of *D. melanogaster* measured in the fall and following the overwintering experiment. Large points are mean average trait values of each time point ± 1 SE. Smaller points jittered behind the means show raw average values for each independent population. A. Dry mass measured in milligrams (mg) between fall and overwintering populations. B. Fecundity measured as average egg production across a three-day assay between fall and overwintering populations. C. Starvation tolerance measured as average time to mortality between fall and overwintering populations. D. Spinosad resistance measured as the percent egg:adult survival on insecticide media.

### Tradeoffs between overwintering and insecticide resistance

To test for trade-offs between overwintering selection and insecticide resistance, we compared phenotypic performance in fitness associated traits between fall and overwintering control and resistant populations. Importantly, three of the five resistant and one of the eight control populations went extinct during the winter and therefore reduced our replication for this contrast.

With only two surviving resistant replicates, we focused on the effect size difference between population types (Hedge’s G: control minus resistant) at both time points and reported these values to show the magnitude of the differences in groups along with associated test statistics and P values (Table S3).

We found that for both body size and fecundity, the effect size between control and resistant populations increased from fall to overwintering. Specifically, there was greater dry mass for overwintering resistant and greater fecundity for overwintering control populations (Dry mass: Figure 4A; control minus resistant Hedge’s G fall = –0.78 [-1.49, –0.05], overwinter = –1.05 [-1.97, –0.12]; Fecundity: Figure 4B; control minus resistant Hedge’s G fall = 0.41 [0.04, 0.078], overwinter = 1.42 [0.85, 1.98]). In both models, time point and the interaction of time point and population type were significant predictors, but there was no main effect of population type (Table S3). We did not, however, find any differences in starvation tolerance (Figure 4C; Table S3; control minus resistant Hedge’s G fall = –0.29 [-0.99, 0.41]; overwinter = –0.37 [-1.25, 0.52]) nor were there differences in chill coma recovery time between population types following the winter period (Figure S3; Table S4; control minus resistant Hedge’s G overwinter = –0.42 [-0.87, 0.04]).

**Figure 4.**
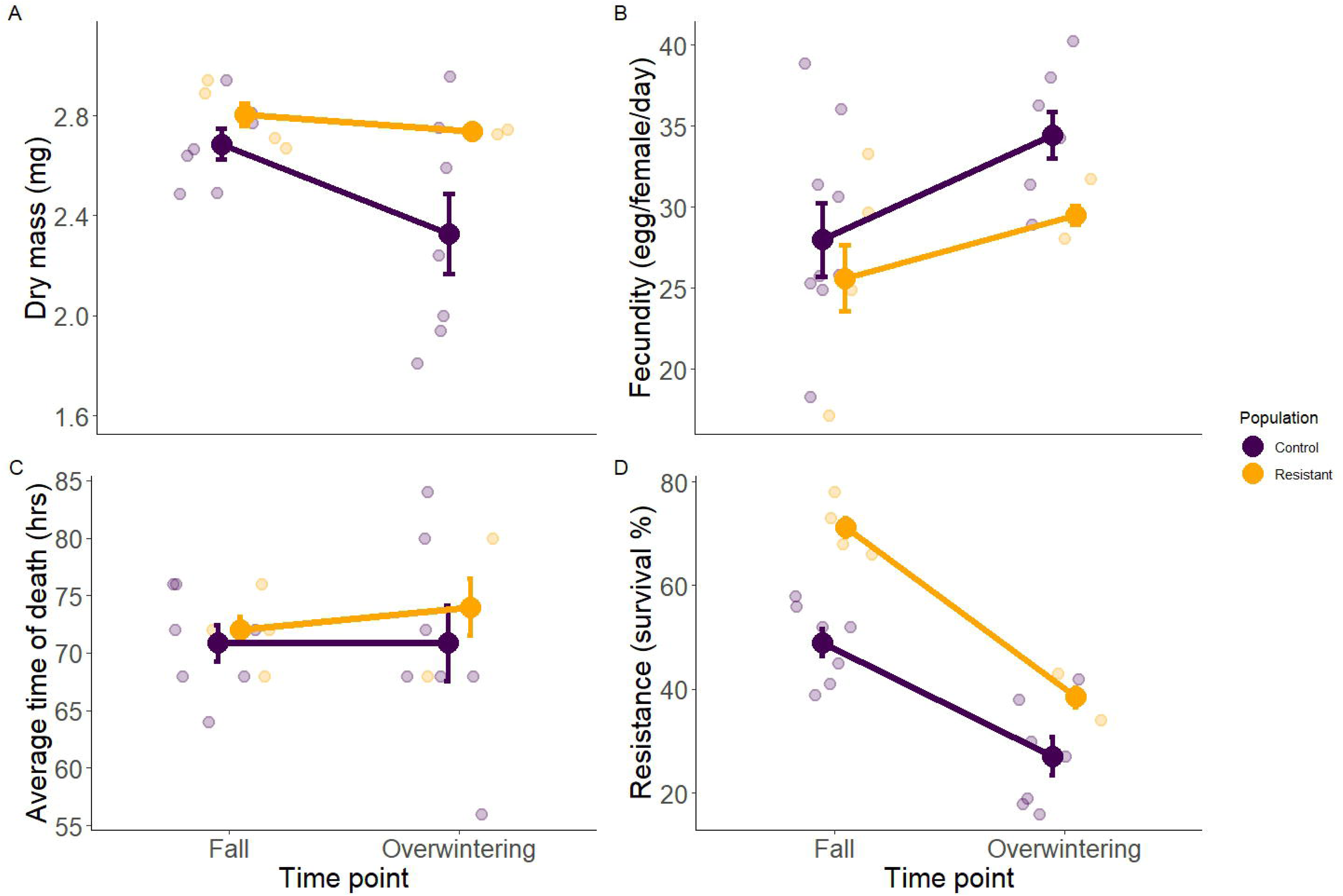
Phenotypic measures of outdoor control populations and outdoor resistant populations of *D. melanogaster* measured in the fall and following the overwintering experiment. Large points show the mean trait value of each population type ± 1 SE. Smaller points jittered behind the means show raw data values for each independent population’s phenotypic measure. A. Average dry mass in milligrams between population types. B. Fecundity measured as average egg production across a three-day assay between fall and overwintering populations. C. Starvation tolerance measured as average time to mortality between fall and overwintering populations. D. Spinosad resistance measured as the percent egg:adult survival on insecticide media between population types.

To specifically test a trade-off in spinosad resistance, we compared egg:adult survival between fall and overwintering time points for both control and resistant populations. We found that the effect size of greater survival between control and resistant population declined in overwintering populations (control minus resistant Hedge’s G fall = –3.52 [-4.10, –2.94], overwinter = –1.90 [-2.30, –1.49]). In our models, there were significant main effects of time point and population type but not for the interaction of time point and population type on resistance (Figure 4D; Table 1).

### Adaptive tracking and overwintering performance

Lastly, we asked if variation in evolutionary responses to winter could be explained by prior evolution in the outdoor environment during the growing season. In our comparison of outdoor reared control and indoor reared founder populations, we did not find any differences in any of the phenotypes (Figure 5; Table S4). In the test for differences in variance between control and founder groups, variance for control populations was significantly greater than founder populations for dry mass, but not for chill coma recovery time, fecundity, or starvation tolerance (Table S5).

**Figure 5.**
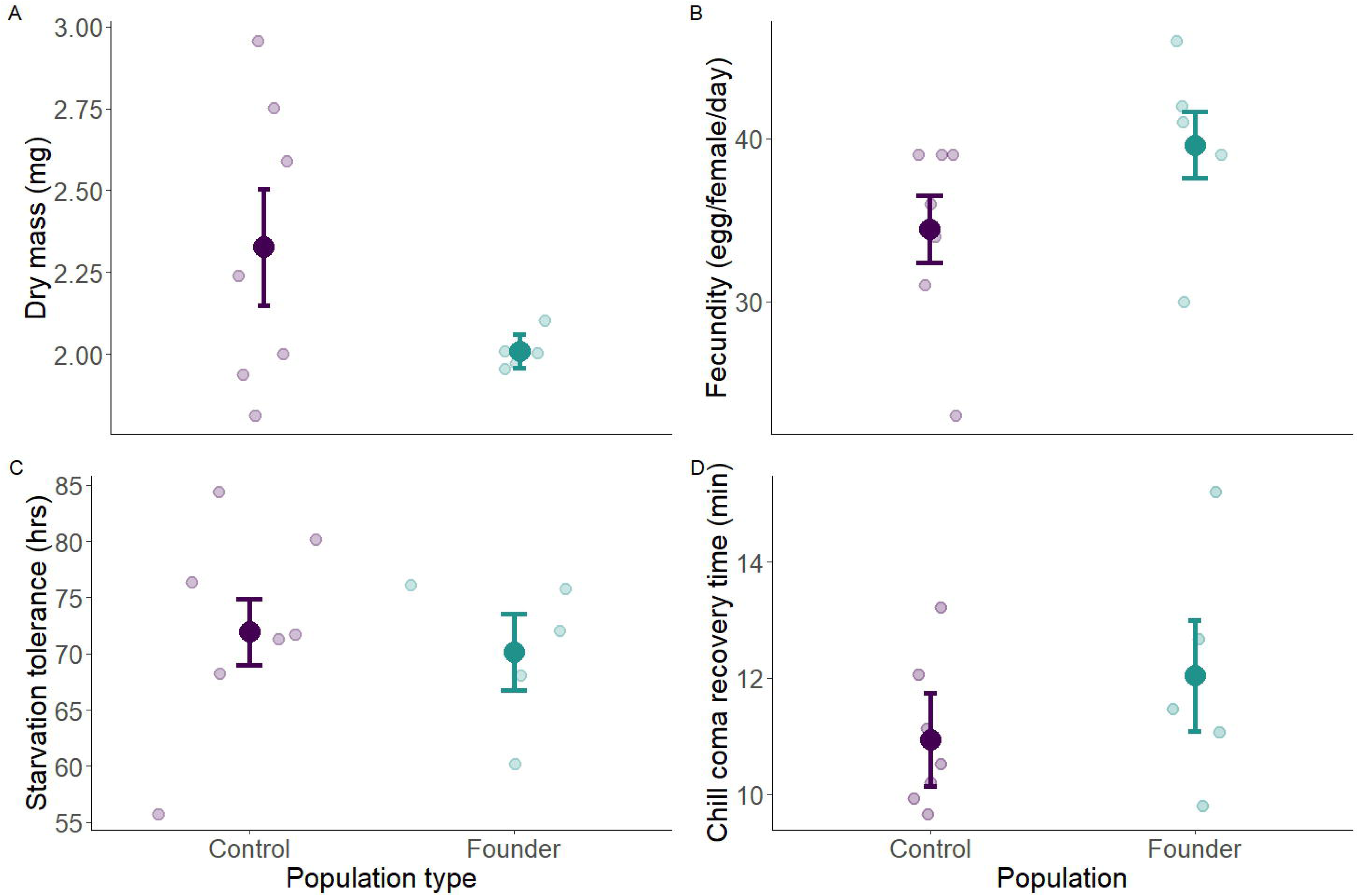
Overwintering phenotypic trait comparison of outdoor control and indoor founder populations. Large points show the mean average trait value of each population type ± 1 SE. Smaller points jittered behind the means show raw data values for each independent population’s phenotypic measure. A. Average dry mass in milligrams between population types. B. Average fecundity across a three-day assay between population types. C. Starvation tolerance measured as the average time of mortality between population types. D. Chill coma recovery time in minutes between population types.

## Discussion

Winter can lead to dramatic declines in temperate insect populations. One potential contributor to population persistence during demographic decline is adaptation, but the nature of the evolutionary trajectories that allow populations to reach high densities the following spring remains poorly understood. Additionally, it is important to understand how insecticide-resistant populations respond to winter selection and if survival negatively covaries with resistance. To these ends, we tested whether overwinter selection results in trait shifts from fall to post-overwintering and if there are trade-offs between previously evolved insecticide resistance and overwintering performance. From overwintering *D. melanogaster* in outdoor mesocosms, we detected evidence for adaptation in multiple phenotypic traits across the winter period, and a putative trade-off between the evolution of insecticide resistance and winter survival. These findings demonstrate that winter conditions can pose strong selective pressures and suggest that this selection may act against spinosad-resistant genotypes. Our findings have important implications for insecticide use in pest management, impacts to non-target species, and overwintering evolution in a changing climate.

### Adaptation across winter

In the outdoor-reared control populations, there was repeated evolution of multiple traits but not always in the predicted direction. From both insect thermal physiology and field data of *Drosophila* (Chown and Nicholson, 2004; James et al. 1997), we expected that overwintering populations would evolve larger body size and reduced fecundity when compared to fall populations, but instead the reverse was true. Previous studies of *D. melanogaster* seasonal dynamics have shown that fall populations exhibit trait evolution towards ‘summer like’ phenotypic values compared to spring populations (Behrman et al. 2015; Machado et al. 2021). One explanation could be that many overwintering *Drosophila* can carry male gametes within their spermatheca from fall matings (Boulétreau-merle and Fouillet, 2002; Collet and Jarman, 2001). Thus, females who successfully overwinter could produce maladapted offspring (Collet and Jarman, 2001).

Another possibility is that body size and fecundity respond in a counter-gradient fashion to the winter period. Body size and reproduction are intimately connected to thermal physiology where larger body size covaries with lower temperature (i.e., Bergmann’s Rule; Partridge et al. 1994; Partridge and Coyne, 1997) and fecundity trades off with stress tolerance and diapause incidence – a pattern found in *D. melanogaster* across latitudinal gradients (Angilletta, 2009; Denlinger and Lee, 2010; Schmidt and Conde, 2006). In a counter-gradient scenario, overwintering selection in high-latitude *D. melanogaster* might favor traits such as greater fecundity, as this could allow populations to jumpstart and access resources more quickly when spring conditions become favorable (Bachmann et al. 2020; Conover et al. 2009).

### Trade-offs between resistance and overwintering

The evolution of resistance during the growing season incurred costs in overwintering performance indicative of a trade-off. First, resistant flies saw greater mortality across winter and second, the effect size of greater spinosad resistance in fall between resistant and control populations decreased overwinter, suggesting that resistant flies that survived the winter period carried reduced resistance. This overwintering susceptibility could be due to physiological costs of spinosad resistance including greater oxidative damage (Weber et al. 2012), minor decreases in longevity, an altered lipid environment, and loss of vision (Perry et al. 2015; Martelli et al. 2022). These detrimental non-lethal effects of resistance could negatively interact with low temperature stress of overwintering leading to the greater mortality we observed. Surprisingly, we did not find the expected differences in cold tolerance and starvation tolerance between resistant and control populations as drivers of lower overwintering survival. We interpret these phenotypic results with caution, however, as these findings might be due to survivorship bias and the reduced sample size for overwintered resistant populations limited the power to detect differences in these traits.

### Adaptive tracking and overwintering phenotypic variation

Prior outdoor evolution in control cages did not significantly impact overwintering phenotypes when compared to indoor founder populations. A putative explanation is that our outbred indoor (i.e., founder) populations still retained standing genetic variation and that responses to selection within a generation was sufficient to allow similar overwintering success as was observed in control populations evolving in outdoor conditions (Barrett et al. 2008). Fluctuating selection for multivoltine insects across the growing season can maintain phenotypic variance through temporal balancing selection (Turelli and Barton, 2004; Bertram and Masel, 2019). Based on this, we predicted that phenotypic variance in control populations would be greater than that in founder populations, but this only occurred for body size. Perhaps ‘summer-like’ features of fall-collected control flies combined with warm temperatures leading up to the time of collection (Figure S2; Machado et al. 2021) led to the similar observed variance.

### Interactions and implications of overwintering, resistance, and pest management

Overall, there is little known about interactions between insecticide resistance and overwintering insect biology, and the evidence to date is mixed. In the Colorado Potato Beetle (*Leptinotarsa decemlineata)*, a major agricultural pest, resistant populations stimulate investment in fat body tissue that increase metabolic fuel and inherit epigenetic effects that elicit general stress responses that increase overwintering success (Lehmann et al. 2014; Brevik et al., 2018; Sinclair et al. 2015). In other studies, however, resistant beetles have exhibited lower overwintering survival due to maladaptive behavioral responses (Ferro et al. 2003; Piiroinen et al. 2013). Antagonistic responses between winter stress and insecticide resistance, as in the case of our overwintering *D. melanogaster,* have also been observed in insecticide resistant green peach aphids (*Myzus persicae*) and the mosquito *Culex pipiens* – which showed lower survival overwinter due to greater low temperature susceptibility and differential success in finding overwintering refugia, respectively (reviewed in Kliot and Ghanim, 2012; Bourguet et al. 2004). Given the limited and mixed evidence among insect taxa, more research – specifically with population level replication – is needed to test whether negative correlations between winter survival and insecticide resistance impact population dynamics of pests.

Trade-offs between overwintering and insecticide resistance have important implications for both rapid adaptation and population dynamics under climate change. High latitude insect populations could show reduced rates of resistance evolution – both in target pest species and non-target insects that might provide valuable ecosystem services. If resistance evolution is strongly temperature dependent, however, climate warming could substantially alter the dynamics of resistance evolution (IPCC, 2023; Easterling et al. 2000; Williams et al. 2015). One area of concern is that relaxed selection from winter could lead to greater population level resistance and range expansions of overwintering pest populations. Indeed, with milder winter conditions, resistant pests have expanded into higher latitudes including the diamondback moth (Ma et al. 2021), multiple tick species (Molaei et al. 2022), and Colorado Potato Beetle (Piiroinen et al. 2013). Of course, this also applies to the evolution of resistance in non-target beneficial insect species (e.g., pollinators) where relaxed selection for low temperatures could lead to an evo-system service (Rudman et al. 2017). However, winter climate change is also associated with greater variability and is hypothesized to lead to population decline (Williams et al. 2015; Sinclair, 2013) or select for generalist phenotypes (Pfennig et al. 2020; Marshall et al. 2020). Given these different outcomes, the capacity for adaptation across winter and trade-offs between resistance and winter survival are important starting points for determining the effects of winter climate change on temperate insects.

### Conclusion

In this study, we add important evolutionary context for the well-studied ecology and physiology of overwintering ectotherms. Notably for *D. melanogaster*, adaptation leading to population persistence (and maintenance of genetic variation) overwinter seem to be critical for the ‘spring reset’ where greater fecundity can allow rapid population growth and recolonization of resources at the start of the next growing season (Machado et al. 2021; Behrman et al. 2015). That winter led to greater mortality in resistant populations demonstrates potential costs to spinosad-resistant flies in their ability to make this same reset. More broadly, this work has implications for understanding seasonal demographics of important agricultural pollinators, management of pest species, and rapid adaptation of ectotherms in temperate climates.

## Supporting information

Supplemental material including additional analyses and results

## Data availability statement

Data are available via the Open Science Framework, https://osf.io/mw9r6/?view_only=817453b8efe6412f8db821535d2d5ca8

