## Supplemental material including additional analyses and results for "Adaptation and trade-offs with insecticide resistance in overwintering Drosophila"

**Supplementary material**

*Details on statistical analyses*

Field survival

To test if the number of surviving adult flies following the overwintering period varied among population types, we constructed a generalized linear mixed effects model fit with a binomial error distribution and a logit-link function. We modeled surviving flies as the response variable, population type as a fixed predictor variable, and cage as a random effect.

Spinosad resistance:

To test the resistance of populations to the insecticide spinosad, we measured the mean survival among populations raised on spinosad media. To test differences in resistance across winter, we used percent survivorship in control populations as the response variable with time point as our predictor variable and cage as a random effect. To test differences among overwintering populations, we used percent survivorship as the response variable with population type as a fixed effect predictor variable and included cage as a random effect. Finally, to test differences in resistance between control and resistant populations across the overwintering period, we included population type and time point as fixed effects along with their interaction and included cage as a random effect. We used a poisson error distribution and log-link function for each of these models.

Starvation tolerance:

To test evolution of starvation tolerance before and after the overwintering period, we used the average time to death of each cage as the response variable, time point as the fixed effect variable, and cage as a random effect. We used a gaussian error distribution to fit this model. Next, to test divergent evolution in starvation tolerance among population types across time points, we used the average time to death as the response variable and included population type and time points as fixed effects along with their interaction and cage as a random effect. For this model, our data was best fit with a gaussian error distribution. To test variation in starvation tolerance between control and founder population types we used the average hour of death per individual fly across all replicates from each cage. In our model we considered the hour at death as the response variable, population type as the fixed effect predictor, and cage as a random effect. We modeled the data using a poisson error distribution with a log-link function.

Fecundity:

We averaged the total number of eggs across the three day assay per cage and compared control populations tested at the fall time point and after the overwintering period. To test for differences in fecundity across time points in our model, we used the average number of eggs laid per female per day as a response variable, time point as a fixed effect and cage as a random effect. For this model, our data was best fit with a gaussian error distribution. Next, to test divergent evolution in fecundity among population types across time points, we used the average integer number of eggs as the response variable and included population type and time points as fixed effects along with their interaction and cage as a random effect. For this model, our data was best fit with a gaussian error distribution. Lastly for our third question comparing control and founder populations, average egg production was our response variable, population type was our fixed effect variable and cage was our random effect. In this model, we used a poisson error distribution in this model with a log-link function. For each of these models, we were unable to meet assumptions of homoscedasticity of residuals, likely due to low sample size of overwintered resistant populations. Our efforts to correct for this using different link functions, transforming the response variable, or adding dispersion formulas did not improve model fits so we retained the original models.

Dry mass:

To test for the evolution of body size in control populations across winter, we used the average dry mass as the response variable, time point as a fixed effect, and cage as a random effect. Next, to test divergent evolution in body size among population types across time points, we used the average dry mass as the response variable and included population type and time points as fixed effects along with their interaction and cage as a random effect. In the test of dry mass between control and founder overwintered populations we considered dry mass as the response variable, population type as the fixed effect predictor, and cage as a random effect. For all models we used a gaussian error distribution.

Chill coma recovery time:

To test the potential for evolutionary divergence in cold tolerance among overwintering population types, we used chill coma recovery time as our response variable, population type as a fixed effect predictor variable, and cage as a random effect. For our model, the data was best captured with a gaussian error distribution. We did not have cold tolerance data from our fall populations, so we were unable to compare chill coma recovery time from before and after the overwintering period.

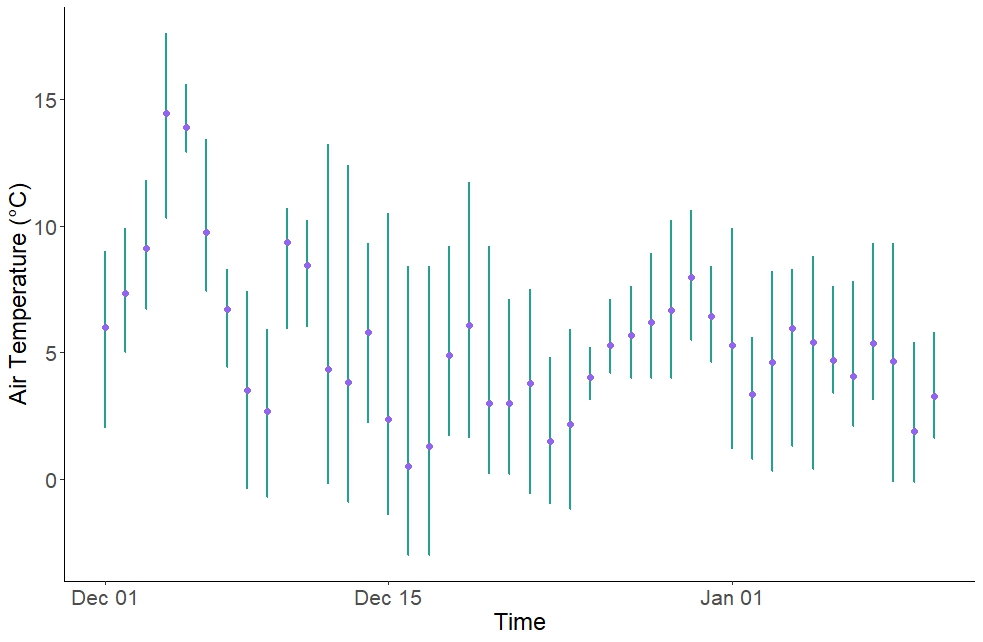

Figure S1. Temperature trace at the Washington State University Vancouver orchard in 2023 - 2024 during the overwintering experiment. Purple points are the mean air temperature in Celsius and the green line displays the full temperature range per day.

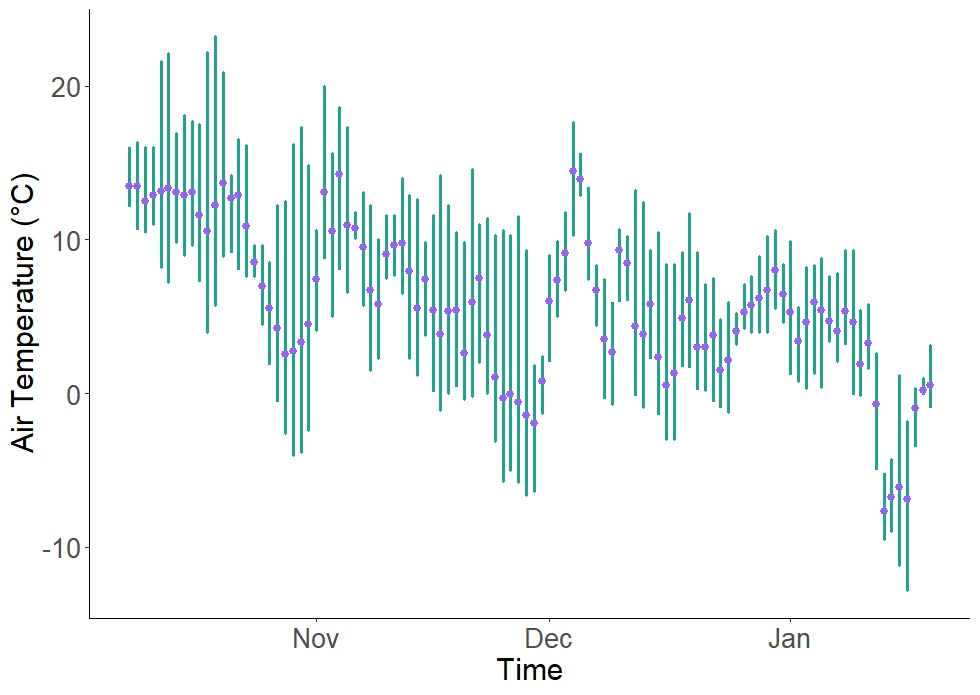

Figure S2. Temperature trace at the Washington State University Vancouver orchard in 2023 - 2024 prior to collection and throughout the overwintering experiment. Purple points are the mean air temperature in Celsius and the green line displays the full temperature range per day. The date of egg collection was October 30th and the overwintering period took place from December 1st through January 10th.

Table S1. Indoor incubator temperature and light-cycle schedule to simulate spring-phenology cues following the overwintering period.

| **Date** | **Diurnal light cycle** | **Temperature** |
| --- | --- | --- |
| 1/10/2025 | 16:8 LD | 15:8 °C |
| 1/12/2025 | 16:8 LD | 18:11 °C |
| 1/15/2025 | 16:8 LD | 20:15 °C |
| 1/19/2025 | 16:8 LD | 25:18 °C |

Table S2. Schedule of phenotyping assays for Drosophila following the overwintering and spring acclimation period.

| **Phenotype measured** | **Steps** | **Date** |
| --- | --- | --- |
| Resistance (survival on spinosad media) | Collected eggs from 2nd common garden adult flies and transferred onto spinosad and control media vials | 2/19/2024 |
| Chill coma recovery time | Transferred females flies to empty centrifuge tube; ran assays | 2/21/2024 |
| Fecundity | Transferred flies to fecundity laying containers | 2/22/2024 |
| Starvation tolerance | Transferred male flies to agar vials | 2/22/2024 |
| Fecundity | Day 1 measure of egg counts | 2/23/2024 |
| Fecundity | Day 2 measure of egg counts | 2/24/2024 |
| Starvation tolerance | Last check of survival | 2/25/2024 |
| Dry mass | Incubation for 24 hours at 55 C | 2/27/2024 |
| Dry mass | Weighed dried flies on microbalance | 2/28/2024 |
| Resistance (survival on spinosad media) | Counted survival of adults | 3/2/2024 |

Table S3. Estimates of model outputs and statistical tests for each phenotypic response from fall and overwintering control and resistant populations with main effect terms of time point and population type and the interaction of both terms.

| **Phenotypic measure** | **Term** | **Estimate** | **SE** | **Χ^2^** | **P** |
| --- | --- | --- | --- | --- | --- |
| Dry mass (mg) |  |  |  |  |  |
|  | Time point | -0.398 | 0.0523 | 57.8 | **< 0.0001** |
|  | Population type | 0.0848 | 0.138 | 0.378 | 0.539 |
|  | Time : Population | 0.327 | 0.104 | 9.93 | **0.00162** |
| Fecundity (egg/female/day) |  |  |  |  |  |
|  | Time point | 5.47 | 1.52 | 120.7 | **< 0.0001** |
|  | Population type | -2.40 | 2.51 | 0.911 | 0.340 |
|  | Time : Population | -2.91 | 1.05 | 7.65 | **0.00568** |
| Starvation tolerance  (avg. hour mortality) |  |  |  |  |  |
|  | Time point | 0.0001 | 1.64 | 0.250 | 0.617 |
|  | Population type | 1.14 | 2.61 | 0.813 | 0.367 |
|  | Time : Population | 2.91 | 3.32 | 0.770 | 0.380 |
| Spinosad resistance  (mean survival) |  |  |  |  |  |
|  | Time point | 21.7 | 1.05 | 174.7 | **< 0.0001** |
|  | Population type | 22.5 | 1.08 | 25.2 | **< 0.0001** |
|  | Time : Population | 0.998 | 1.09 | 0.000800 | 0.980 |

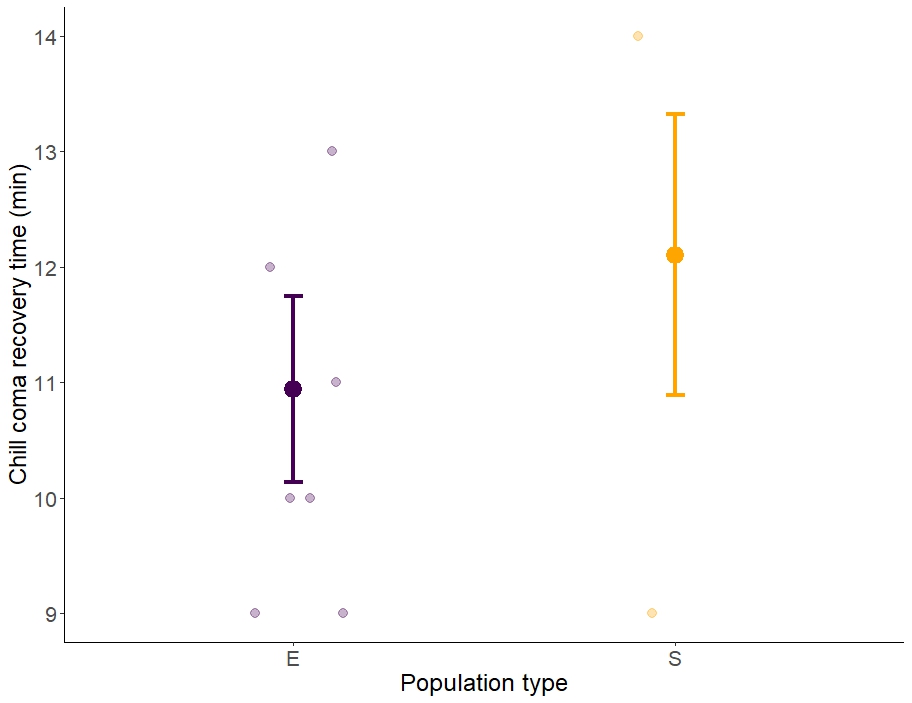

Figure S3. Plot of chill coma recovery time for resistant (S) and control (control) fly populations. Large points show the mean trait value of each population type ± 1 SE. Smaller points jittered behind the means show raw data values for each independent population’s phenotypic measure.

Table S4. Model output of estimate for chill coma recovery time between resistant and control populations just in post-overwintered populations.

| **Trait** | **Term** | **Estimate** | **SE** | **Χ^2^** | **P** |
| --- | --- | --- | --- | --- | --- |
| Chill coma recovery time |  |  |  |  |  |
|  | Population type | 1.21 | 1.21 | 1.01 | 0.316 |

Table S5. Model estimates and results for tests of overwintering outdoor control and indoor founder populations.

| **Trait** | **Term** | **Estimate** | **SE** | **Χ^2^** | **P** |
| --- | --- | --- | --- | --- | --- |
| Dry mass (mg) |  |  |  |  |  |
|  | Population type | 0.409 | 0.291 | 0.198 | 1.59 |
| Fecundity (average egg production) |  |  |  |  |  |
|  | Population type | 0.143 | 0.0937 | 2.33 | 0.127 |
| Chill coma recovery time (minutes) |  |  |  |  |  |
|  | Population type | 1.08 | 0.869 | 1.54 | 0.215 |
| Starvation tolerance (mean average time to death, hours) |  |  |  |  |  |
|  | Population type | -0.0253 | 0.0638 | 0.157 | 0.692 |

Table S6. Levene’s test of differences of variance between control and founder populations for each phenotype.

| **Phenotype** | **Variance control** | **Variance F** | **Levene’s Test** | **P** |
| --- | --- | --- | --- | --- |
| Dry mass | 0.222 | 0.0182 | F_1,10_ = 10.0 | **0.01** |
| Fecundity | 10402 | 11900 | F_1,10_ = 0.0400 | 0.850 |
| Chill coma recovery time | 4.52 | 6.42 | F_1,10_ = 0.450 | 0.520 |
| Starvation tolerance | 71.6 | 36.1 | F_1,10_ = 0.190 | 0.670 |
